# Inclusion of glycopeptides in hydrogen/deuterium exchange mass spectrometry analysis of SARS-CoV-2 spike ectodomain provides in-creased sequence coverage

**DOI:** 10.1101/2023.06.14.544985

**Authors:** Christopher A. Haynes, Theodore R. Keppel, Betlehem Mekonnen, Sarah H. Osman, Yu Zhou, Adrian R. Woolfitt, Jakub Baudys, John R. Barr, Dongxia Wang

## Abstract

Hydrogen/deuterium exchange mass spectrometry (HDX-MS) can provide precise analysis of a protein’s conformational dynamics across varied states, such as heat-denatured vs. native protein structures, localizing regions that are specifically affected by such conditional changes. Maximizing protein sequence coverage provides high confidence that regions of interest were located by HDX-MS, but one challenge for complete sequence coverage is N-glycosylation sites. The deuteration of glycopeptides has not always been identified in previous reports of HDX-MS analyses, causing significant sequence coverage gaps in heavily glycosylated proteins and uncertainty in structural dynamics in many regions throughout a glycoprotein. We report HDX-MS analysis of the SARS-CoV-2 spike protein ectodomain in its trimeric pre-fusion form, which has 22 predicted N-glycosylation sites per monomer, with and without heat treatment. We identified glycopeptides and calculated their isotopic mass shifts from deuteration. Inclusion of the deuterated glycopeptides increased sequence coverage of spike ectodomain from 76% to 84%, demonstrated that glycopeptides had been deuterated, and improved confidence in results localizing structural re-arrangements. Inclusion of deuterated glycopeptides improves the analysis of the conformational dynamics of glycoproteins such as viral surface antigens and cellular receptors.

**Abstract Figure:** 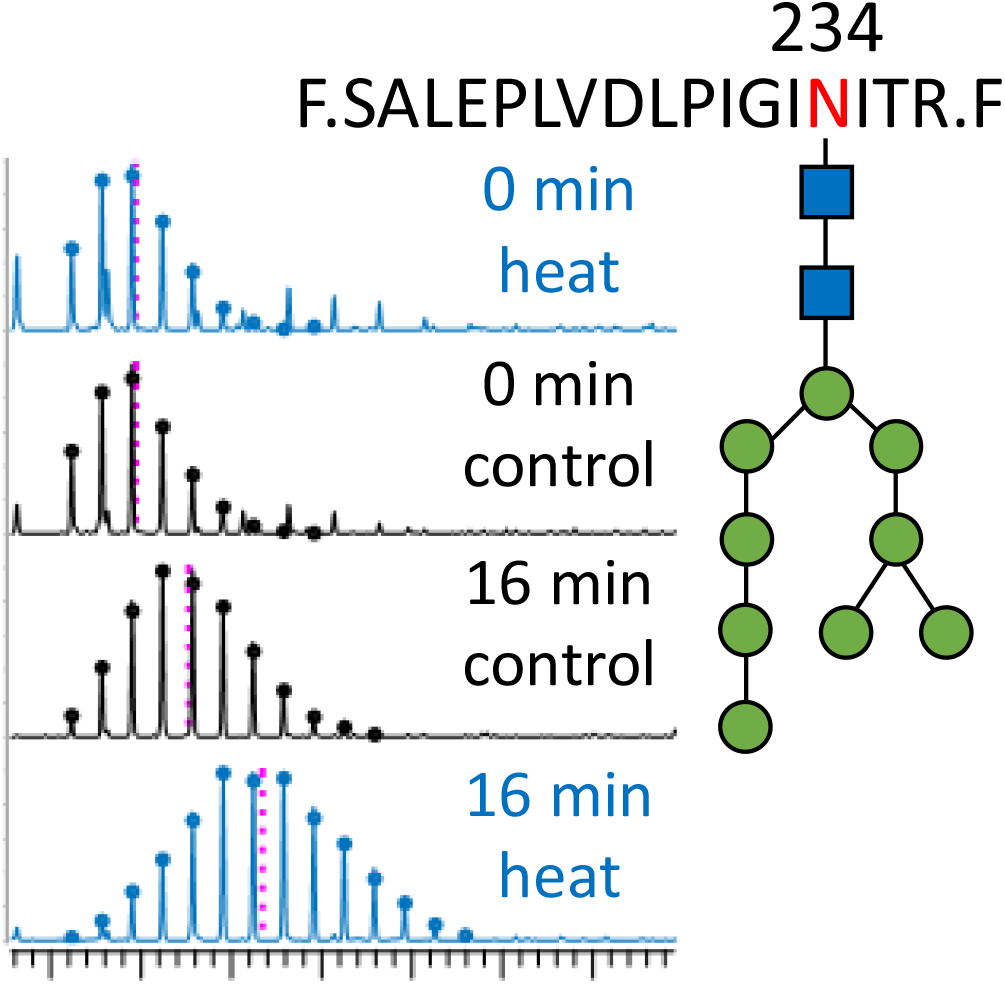

## INTRODUCTION

Hydrogen/deuterium exchange mass spectrometry (HDX-MS) provides a framework of sample preparation, proteolytic digestion, peptide identification, and deuterium uptake measurement over time to identify changes in protein conformation (1, 2). Peptides with changes in amide backbone hydrogen exchange typically indicate movement or stabilization of alpha helices, beta sheets, and other hydrogen bonds contributing to secondary structure (3). Solvent accessibility also plays a role in deuterium labeling of proteins (4). One goal of HDX-MS studies is maximizing protein sequence coverage, since gaps could include regions with informative deuterium labeling. For example, localization of a monoclonal antibody’s epitope (5) can be challenging if sequence coverage of the antigenic protein is incomplete (6, 7, 8).

Sequence coverage gaps in HDX-MS experiments are often associated with an N-glycosylation sequon (NXS/T, where X is not Pro) and attached glycan moiety. Detecting and identifying glycopeptides and their glycans requires advanced tandem mass spectrometry (MS2) methods and appropriate data processing for detection and assignment of glycopeptides (9, 10, 11, 12). We have now combined recent advances in instrumentation and software to enable combining glycopeptide proteomics and HDX-MS to allow the inclusion of N-glycosylated peptides to improve sequence coverage.

SARS-CoV-2 spike, a surface glycoprotein of the virus causing COVID-19 (13, 14), has 22 potential sites of N-glycosylation per monomer (15, 16). The high level of glycosylation has caused significant gaps in sequence coverage and incomplete HDX-MS data (13, 17, 18). Spike protein is the surface antigen and target of all currently available COVID-19 vaccines (19). Its binding to angiotensin-converting enzyme 2 (ACE2) is critical to penetration of a host cell and initiation of viral infection (17, 20, 21). During our HDX-MS analyses of spike we have now applied a previously described method for detecting glycopeptides (22, 23) to the deuterium-labeled D614G variant (24). Heat-treatment of spike was used to significantly change protein structure and demonstrate the utility of deuterated glycopeptide data to improve HDX-MS conformational analysis.

## EXPERIMENTAL SECTION

### Chemicals and Reagents

Phosphate-buffered saline (PBS) tablets, LC-MS grade water 0.1% formic acid, methanol and acetonitrile 0.1% formic acid were from Fisher Scientific (PA). Deuterium oxide (99.9%) was from Cambridge Isotope Laboratories (MA). Urea and tris(carboxyethyl)phosphine (TCEP) were from Sigma-Aldrich (MO). Reagent and sample vials for HDX-MS were from Trajan Scientific and Medical (NC) and Thermo Scientific (CA), respectively.

### Plasmid Design and Cell Culture

The SARS-CoV-2 D614G spike protein ectodomain expression plasmid was designed as follows: amino acids 1-1208, furin cut site PRRAR substituted to PGSAS, 6 HexaPro stabilization substitutions (K986P, V987P, F817P, A892P, A899P, A924P), a T4 fold self-trimerization domain, and a polyhistidine tag for purification by nickel-charged nitrilotriacetic acid (Ni-NTA). FASTA sequences for 614G and Omicron spike constructs are in Supporting Information. Expi293F cells were obtained from ThermoFisher as part of the Expi293F protein expression system. Cells were cultured for 3+ passages at 37°C/125 RPM/8% CO_2_ not exceeding a density of 5 × 10^6^ cells/ml or passage >30, seeded at a density of 2.5 × 10^6^ cells/mL one day prior to transfection, and diluted to 3 × 10^6^ cells/mL immediately before transfection. Plasmid was transfected at 1 μg DNA per 1 mL of cell culture using ExpiFectamine and Opti-Mem I Reduced Serum Medium (Thermo Scientific). Transfected culture was left to express for four days at 30°C/125 RPM/8% CO_2_ before harvesting and centrifuging twice at 4°C/3000 RCF for 10 minutes to clarify the supernatant.

### Protein Purification

Three mL of 1 M imidazole (Sigma-Aldrich) per 100 mL of transfected clarified supernatant was added prior to purification. Purification used two buffers, one binding buffer (20 mM sodium phosphate, 500 mM sodium chloride, 30 mM imidazole pH 7.4) and one elution buffer (20 mM sodium phosphate, 500 mM sodium chloride, 500 mM imidazole pH 7.4). The column was first equilibrated with 5 column volumes (CV) of binding buffer. The supernatant was then loaded onto an AKTA Avant 150 FPLC through a Cytiva 5 mL HisTrap FF column at 0.5 mL/min. After sample loading, the column was washed with 5 CV binding buffer. The protein was eluted off the column using a gradient elution from 100% binding buffer to 100% elution buffer over 5 CV and peaks were collected in 2 mL fractions at 4°C. The peak fractions were loaded onto a 7K MWCO Zeba desalting column at 4°C, buffer exchanged into 300 mM sodium chloride, 20 mM Tris-HCl pH 8, and later concentrated on an Amicon 100K MWCO Ultra-15 centrifugal filter column at 4000 RCF at 4°C until desired concentration/volume was met. A minimum of 500 μg of this affinity purified protein was then loaded onto a Cytiva Superose 6 Increase 10/300 gel filtration column also on the AKTA Avant 150 system. Preparative size exclusion chromatography took place at 0.5 mL/min in 300 mM sodium chloride, 20 mM Tris-HCl pH 8. Peaks corresponding to the size of spike trimer were then fractioned into 2 mL fractions and later concentrated on a 100K MWCO Amicon filter if needed. Protein quality was checked by SDS-PAGE gel and analytical size exclusion chromatography before use.

### Heat-treatment, Trypsin Digestion and Gel Analysis of Spike Ectodomain

Spike ectodomain protein (Omicron variant, 1.3 μg/μL in 20 mM Tris, 300 mM NaCl, pH 8) was diluted 1:20 in HA buffer (100 mM Tris, 50 mM NaCl, pH 7.8) and 40 μL aliquots were incubated in a PCR tube and heater (Thermo Scientific) for 1 hour at 25, 35, 45, 55, and 65°C. After cooling to room temperature and centrifugation (1000 RPM, 10 min), supernatants were transferred to clean PCR tubes for trypsin digestion. Sequencing grade trypsin (Promega, WI) was diluted in HA buffer so that adding 2 μL of trypsin to each heat denatured aliquot resulted in a 1:100 molar ratio of trypsin to spike. A control set of aliquots had 2 μL of HA buffer with no trypsin added to verify heat-treatment did not result in aggregation. All tubes were incubated for 1 hour at 37°C, cooled to room temperature, measured for volume loss and compensated with HPLC-grade water. A non-reducing Bis-Tris 4-12% gradient gel (Thermo Scientific) was used to detect size-resolved spike proteins (Figure S1).

### Hydrogen/Deuterium Exchange Mass Spectrometry

A DHR-PAL system (Trajan) was used for sample preparation, with sample tray at 20°C, quench tray at 4°C, valve chamber & pre-chiller at 4°C, and digestion chamber at 8°C. Purified spike ectodomain (0.4 μg/μL) was mixed 1:5 (v/v) with PBS prepared using H_2_O (equilibration buffer, measured pH 7.22) or D_2_O (labeling buffer, measured pH 7.56). Labeling times were 0, 60, 240, and 960 sec with 3 technical replicates at each time. Samples were quenched with an equal volume of 2 M urea 0.5 M TCEP pH 2.5 and held at 4°C for 2 min. An UltiMate3000 UPLC system (binary nano pump and loading pump, Thermo Scientific) was used for subsequent online sample handling. Automated valve switching passed the quenched sample over a 2.1 × 20 mm Nepenthesin-2 / Pepsin mixed digestion column (AffiPro, CZ) at 100 μL/min H_2_O 0.1% HCOOH for 2 minutes, trapping the resulting peptides on a 2.1 × 5 mm Fully Porous C18 guard column (Phenomenex, CA), then desalted peptides at 300 μL/min for 4 minutes. Peptides were eluted and resolved by a gradient from 13 to 65% mobile phase B (95:5:0.1 CH_3_CN / H_2_O / HCOOH over 23 min on a 1 × 100 mm Luna Omega 1.6 μm 100 Å C18 column (Phenomenex, CA). A Tribrid Eclipse Orbitrap (OT) mass spectrometer (Thermo Scientific) with HESI-2 electrospray ion source and high-flow needle was operated in positive ion mode to detect peptides for 31 minutes. In all samples precursor scans of resolution 120K (at *m/z* 200) in the range 375-2000 *m/z* were acquired. For each MS1 scan, the top-10 abundant precursor features were selected for data-dependent MS2 scans, selecting for an intensity threshold of 30K counts, monoisotopic peptide precursors, charge states 2^+^ to 8^+^ with an isolation window of 1.2 *m/z*, and not repeating precursor ions more than twice within 15 sec. Precursors were fragmented with HCD at 28% normalized collision energy (NCE) and centroid scanned in the OT with standard automatic gain control (AGC) target, automatic injection time, scan range 120-2000 *m/z*, and resolution 30K. If at least one of three selected oxonium ions was detected (HexNAc 204.0867, HexNAc fragment 138.0545, or HexNAcHex 366.1396) with 15 ppm mass tolerance then EThcD OT-MS2 scans were acquired of that precursor ion. Supplemental HCD was at 20% NCE with profile scans from 150-2000 *m/z* at resolution 50K using custom AGC target (500%) and fill time (90 msec). At the end of the analytical gradient, solvent transitioned to 90% mobile phase B for 6 min, and halfway through that time the analytical and trapping columns were put into back-flow washing mode by automated valve switching. After re-equilibration of the analytical column at 13% mobile phase B the injection cycle ended at 45 min. As recommended (25), the entire batch of control and heat denatured samples (all time points and replicates) was randomized for HDX-MS acquisition to minimize batch effects on interpreted differences in protein state, labeling time, and technical replicates. To allow access to the HDX data of this study, the HDX data summary table (Table S2) and the HDX data table (Table S3) are included in the supporting information as per consensus guidelines (25).

### Data Processing

Protein Metrics Inc. (CA) Byos HDX 4.6-37 searched the 6 data files from equilibration buffer samples (0 sec labeling time, control and heat denatured, 3 technical replicates each) to identify (glyco)peptides using MS2 spectra. The search database included spike and both proteases. Both HCD and EThcD tandem mass spectra contributed to peptide spectral matching. Putative (glyco)peptides were then searched for in data files from all samples at the MS level (and appropriate retention times) to identify both unlabeled and deuterated peptides and visualize their isotopic envelopes. Initial spike results (heat-treatment vs. control, 1271 peptides) were narrowed by 1) default software filters (MS2 score > 15, minimum alt_rank_score/primary_rank_score > 0.99, maximum precursor *m/z* error ± 40 ppm, maximum retention time deviation ± 5 min) leaving 1056 peptides, and 2) removing peptides with MS2 score < 150 leaving 106 of 205 glycopeptides, removing peptides with more than ± 10% average, maximum, or minimum “deuteration” in 0 sec samples, and removing peptides causing standard deviations >10% at any labeled time-point, leaving 561 peptides. Additional manual curation involved adjustment of the extracted ion chromatogram (XIC) window used to integrate MS data and generate an isotopic envelope, optimizing the intensity and specificity of that envelope. Peptides with inadequate intensity XICs to estimate deuteration were discarded.

## RESULTS AND DISCUSSION

Traditional HDX-MS analysis of a glycoprotein’s conformational dynamics often leaves large information gaps because of the difficulty in analyzing the deuterium uptake of glycopeptides. During our traditional HDX-MS analysis of SARS-CoV-2 spike protein, we found significant gaps in sequence coverage and a lack of information in many areas of interest (Figure S2). To fill these gaps, we incorporated signature ion-triggered EThcD into HDX-MS analysis.

Identification of glycopeptides used Fourier-transform mass spectrometry (FT-MS) scans to detect precursor ions, and those above an intensity threshold were selected for high-energy collisional dissociation (HCD) FT-MS2 scans. This relatively energetic fragmentation yields abundant b- and y-ions for spectral matching to peptide sequences, but can also produce oxonium ions (such as HexNAc *m/z* 204.0867, HexNAc fragment *m/z* 138.0545, and Hex(2)NAc *m/z* 366.1396) diagnostic for the presence of complex glycosylation of the precursor peptide (10, 22, 26, 27). The presence of these signature oxonium ions triggered data-dependent electron transfer-low energy collision induced-dissociation (EThcD) FT-MS2 of a fresh precursor ion packet, yielding peptide fragments retaining 1 or more hexose groups or intact glycan, ideally confirming the glycosylation sequon’s location (28).

### Glycopeptide inclusion improves sequence coverage

Addition of EThcD to facilitate the detection and identification of glycopeptides during HDX-MS significantly improved sequence coverage and conformational dynamic analysis of SARS-CoV-2 614G spike protein. Glycopeptides were confidently identified at 9 out of the 22 potential occupied N-glycosylation sequons per monomer in spike 614G variant (Table S1, 41% sequon coverage and 84% overall coverage) after manual data processing. At all of the 9 N-glycosylation sequons with peptide coverage only glycopeptides were identified (with a single exception, N1074 was covered by one non-glycosylated peptide). Therefore, without the inclusion of EThcD for glycopeptide detection and identification, the sequon coverage would have dropped to 1 (5%) and overall spike coverage to 76% (Figure S2). Additionally, because nine N1074 glycopeptides had HexNAc(2)Hex(5) and HexNAc(2)Hex(6) glycans, the detection of the unfilled, non-glycosylated peptide may not represent the true conformational dynamics at this site.

Sequon-specific combinations of peptide and glycan diversity were observed. Seven glycopeptides (one in 2^+^ and 3^+^ charge states) with unique amino acid sequences including N234 displayed only one glycan type HexNAc(2)Hex(9). In contrast, 13 unique glycopeptide amino acid sequences including N603 displayed five or more glycan types, while the only glycopeptide amino acid sequence including N1134 displayed ten glycan types. Table S1 summarizes sequon coverage and occupancy. All glycan groups at the 9 sequons with coverage were of similar types (oligomannose, complex, and hybrid) as described in previous publications (11, 22, 29).

Evidence for deuterated glycopeptides is shown in Figures 1 to 3 using glycopeptide 597-607 HexNAc(2)Hex(5) as an example. We obtained high-confidence (Byonic score = 508.5) MS2 identification (Figure 1), reproducible extracted ion chromatograms (Figure 2A), and high-quality isotopic envelopes (Figure 2B). We observed that each unique amino acid sequence for a glycopeptide showed consistent uptake plots in both control and heat denatured states although they had different N-glycan groups. For example, glycopeptides with sequence ^597^VITPGTNTSNQ^607^ had highly consistent deuterium uptake (Figure 3, left peptide) even though 5 different glycan groups were identified on N603. However, overlay of uptake plots for twelve N603 overlapping glycopeptides (Figure 3) indicated that unique amino acid sequences had different uptakes, and increasing glycopeptide length affected estimated deuteration. For example, C-terminally extending the above glycopeptide by 2 amino acids decreased uptake in the control state but increased uptake after heat-treatment (Figure 3, center peptide). Our interpretation was that glycopeptide amino acid sequence was a stronger determinant of deuteration than N-glycan identity.

**Figure 1.**
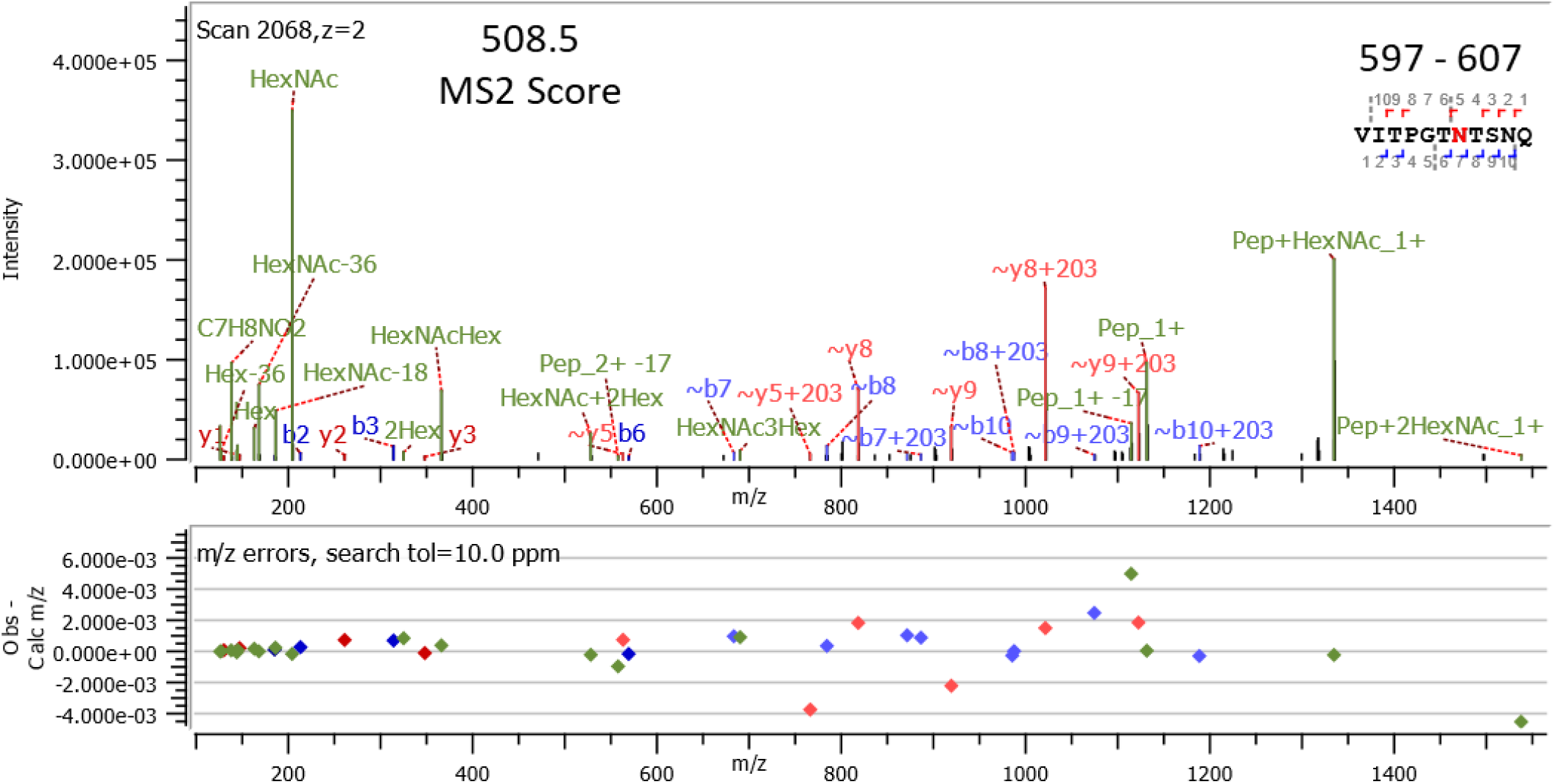
Tandem mass spectrum of glycopeptide 597-607 HexNAc(2)Hex(5) including sequon N603. Sample was unlabeled (0 sec D_2_O exposure), fragment mass errors are shown in the lower panel.

**Figure 2.**
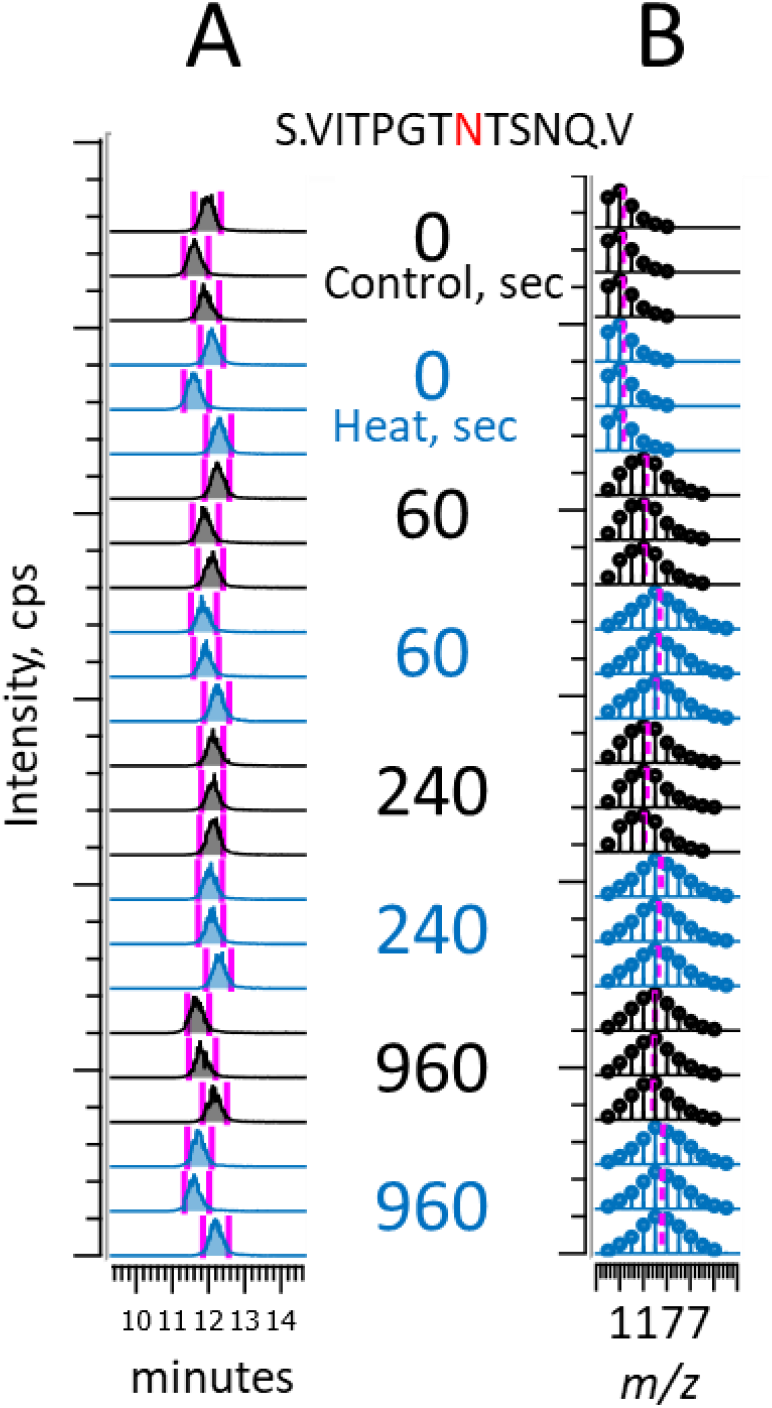
Extracted ion chromatograms (A) and isotopic envelopes (B) of glycopeptide 597-607 HexNAc(2)Hex(5) including sequon N603 (red N). Technical triplicate injections at each D_2_O exposure time (central numbers) and both control (black) and heat treatment (blue) conditions were injected in randomized order.

**Figure 3.**
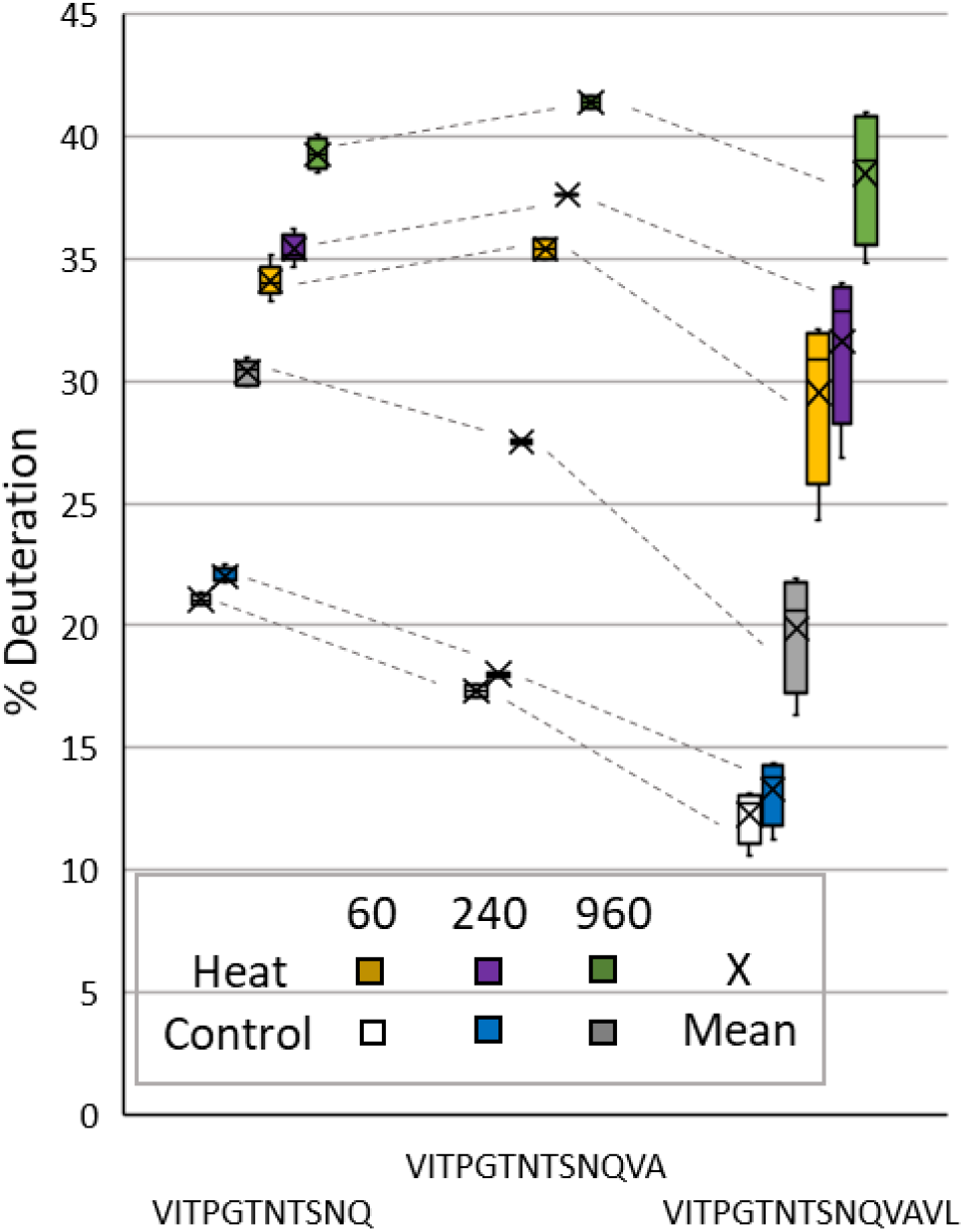
Box and whiskers uptake plots of 12 glycopeptides including sequon N603. The left, center, and right boxes are N = 6, 2, and 4 unique glycopeptides (sequences, glycans, or charge states), respectively. Dashed lines linking boxes indicate the same D_2_O exposure time and control or heat denatured state.

Supporting this interpretation, glycopeptides covering sequons N61 and N234 showed two levels of deuteration in the heat denatured state on over-lay uptake plots, also associated with the length of glycopeptides. For example, the N61 sequon’s proximity to the sequence ^66^HAIH^69^ indicated that inclusion of this histidine-rich “cool” spot of deu-teration decreased the percentage labeling of 7 longer glycopeptides significantly compared to 4 overlapping but shorter glycopeptides that did not include the ^66^HAIH^69^ sequence (Figure S3). This result is consistent with the slow deuteration of His residues (30, 31).

### Isotopologue ratios of deuterated glycopeptide fragment ions

Glycopeptide deuteration (Figures 1 to 3) and the presence of an amide bond in N-acetyl-hexosamines suggested deuteration was also being measured in the N-acetyl amide group(s) of N-linked glycans, which has been previously confirmed at the glycan and glycopeptide levels (32) and recently reviewed (33). To confirm this labeling, isotopologue ratios of glycopeptide ^597^VITPGT<u>N</u>TSNQ^607^ HexNAc(2)Hex(5) fragment ions were compared for both control and heat denatured states at all deuteration times. All MS2 scans consistent with this glycopeptide precursor’s *m/z* and LC retention time were converted to lists of product ion *m/z* vs. intensity, and a ratio was taken between a fragment’s monoisotopic *m/z* (M0) and its M + 1 isotope’s *m/z* (M1, Figure 4 *inset*)). First, the theoretical M0/M1 ratio for HexNAc ion ([C_8_H_14_NO_5_]^+^, M0 *m/z* 204.0867) is 11.5, similar to the observed 9.6 ± 2.9 (Figure 4, C0 and H0, white bars). Second, the theoretical M0/M1 ratio for ∼y8 ion ([C_31_H_52_N_11_O_15_]^+^, fragment ^600^PGT<u>N</u>TSNQ^607^ without the glycan attached, Figure 1) is 2.99, similar to the observed 2.24 ± 1.21 (Figure 4, C0 and H0, grey bars). Third, the ∼y8 ion’s M0/M1 ratio (Figure 4, grey bars) significantly decreased after D_2_O exposure, consistent with deuteration of this peptide’s 7 amide groups, reducing the amount of M0 *m/z* 818.3639 and increasing the amount of M1 *m/z* 819.3779. Fourth, the HexNAc ion’s M0/M1 ratio (Figure 4, white bars) also decreased after D_2_O exposure, indicating that it was labeling during the 16 minutes of this experiment. Student’s T-test of the HexNAc M0/M1 ratios at 0 and 960 seconds of D_2_O exposure indicated a significant difference (p = 0.002), so these results collectively support deuteration at the amide proton in the N-acetyl moieties on glycans N-linked to glycopeptides. Similar data for a different glycopeptide, 1132-1145 HexNAc(4)Hex(3)Fuc(1) is shown in Figure S4.

**Figure 4.**
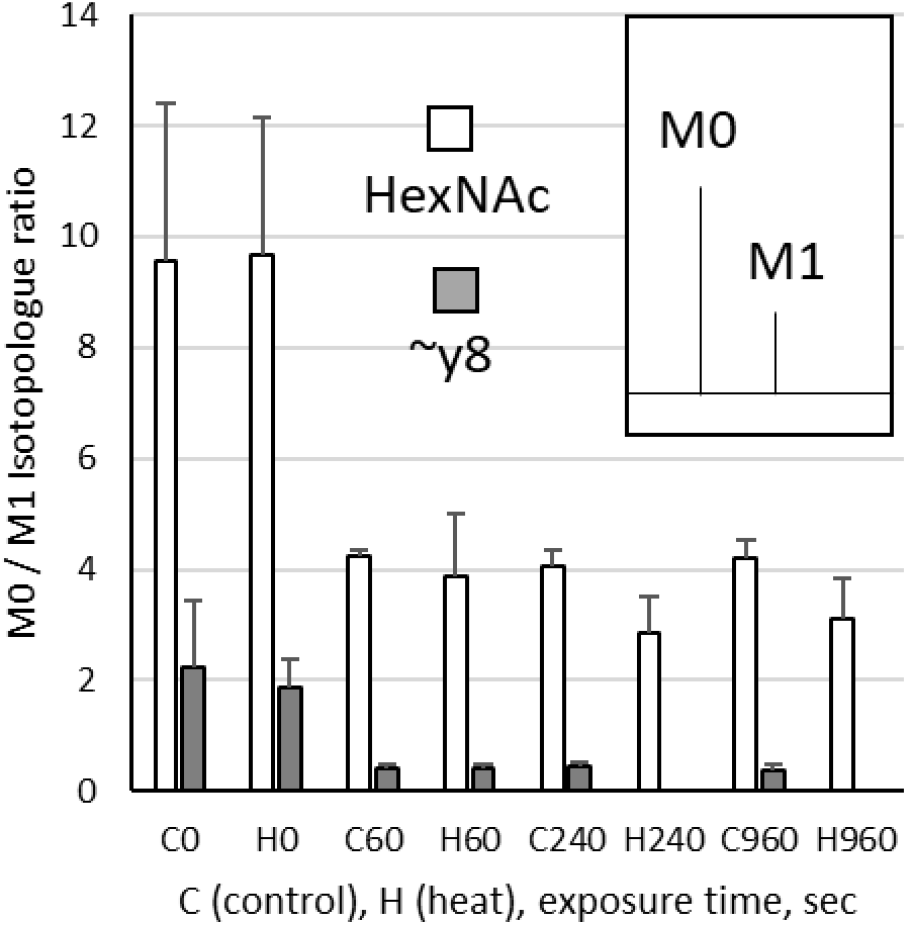
Isotopologue ratios of glycopeptide 597-607 HexNAc(2)Hex(5) fragment ions (see Figure 1 for a typical MS2 spectrum of this peptide). *Inset*: the ratio of M0 (monoisotopic peak) to M1 (first M + 1 isotopic peak) was measured for HexNAc (204/205) and ∼y8 (818/819) in a representative technical replicate at each state and D_2_O exposure time. As fragment ions shifted to a deuterated distribution the ratio decreased. N = 3 to 6 averaged MS2 scans for each bar, error bars are one standard deviation. At H240 and H960 the ∼y8 M0 ion was not detected.

Deuteration of glycans is significant because calculation of “maximum peptide deuteration” is based on the number of amino acid amides that could have been labeled (all except the N-terminal 1 or 2 residues and any proline residues (34, 35)), so the presence of each N-acetyl moiety in a glycan structure could add one potential labeling site per glycopeptide. Additional complexity is added by the variability of glycans at a given N-glycosylation sequon. We will continue to explore the maximum peptide deuteration level for different types of glycopeptides.

### Control state: no heat-treatment

The workflow to include glycopeptides in HDX-MS analysis was then tested by comparing native and heat denatured spike protein. Without heat treatment, spike ectodomain (glyco)peptides showed a range of HDX labeling (structural dynamics) between ∼0% and ∼35% based on sub-domain location (Figure 5A). Localization of these patterns on a model based upon Protein Data Bank (PDB) files 6VSB and 6VXX (36) included key domains (furin cleavage site, fusion peptide, 630 loop, etc.) often missing from RCSB structures (www.rcsb.org), possibly because disordered regions of spike can have low electron density in cryo-EM analysis (37). The least deuterated area (<5%) was the trimeric core interface (764-782 helix, 997-1020 helix, and 1043-1062 strand in β-sheet), as previously described (17, 18).

**Figure 5.**
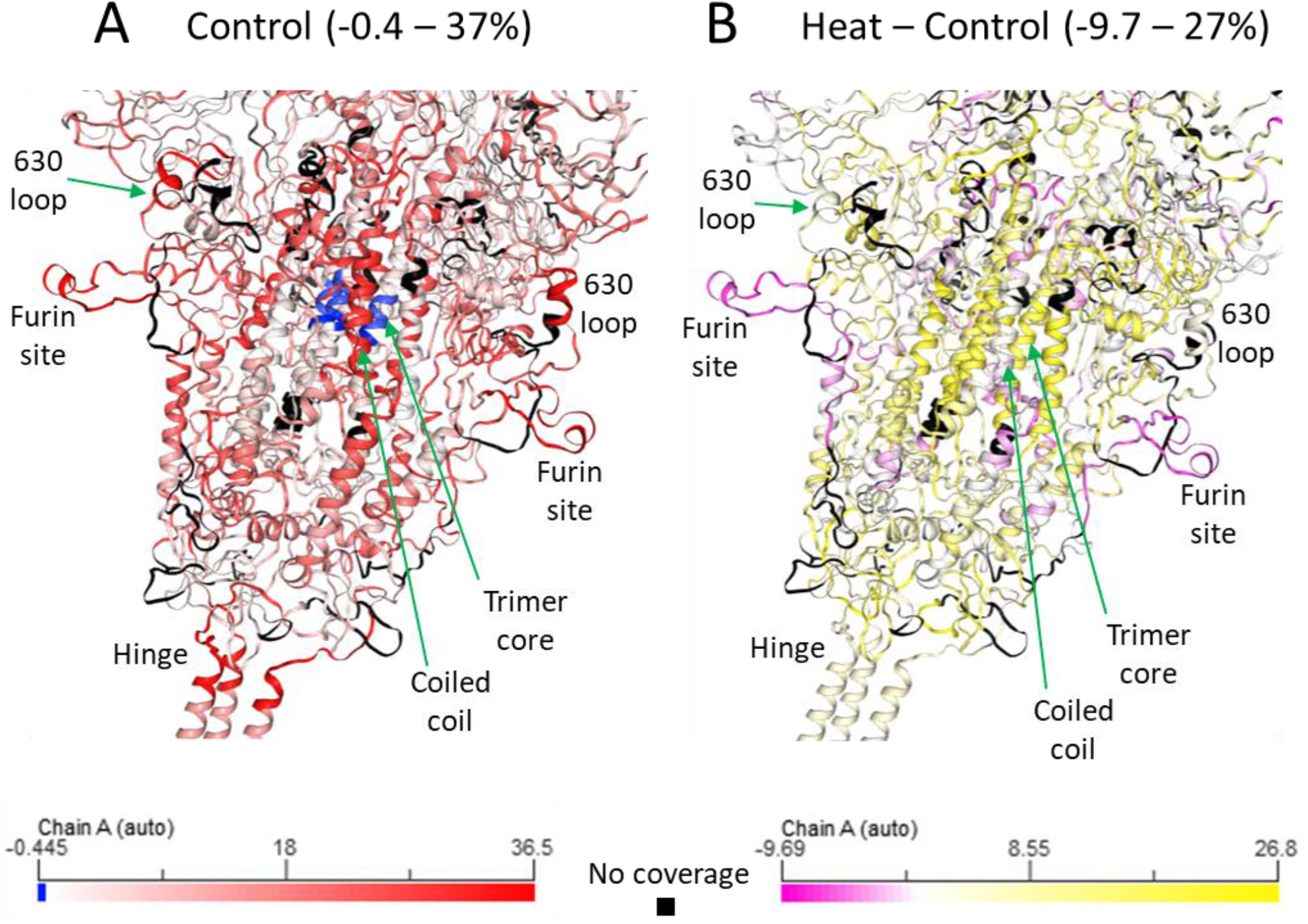
Heat map data (960 sec) overlaid on a model based upon PDBs 6VSB and 6VXX (36), view of central trimer head with stalk at lower left corner. Note the furin cleavage site labeled 35% in the control state (A) is labeled -8% in the (heat – control) visualization (B).

The most labeled regions (35%) were the “hinge” between head and stalk regions (1132-1145), furin cleavage site (672-690), and 630 loop (624-636), also consistent with a previous report (18). Labeling of the “hinge” is a novel HDX-MS result, completely based on the coverage of deuterated glycopeptides containing glycan-modified N1134 (Figures 5 and S2). Robust labeling (24-30%) was observed in the fusion peptide (823-851), N-terminus (19-31), and middle helix of the stalk (1174-1197).

The receptor binding domain (RBD) had the highest deuteration (27%) on a strand forming its hinge with the rest of S1 (319-346, including glycopeptide coverage) and the two loops of the receptor binding motif (RBM, 442-452 and 471-487) for ACE2 (38). The other strand forming the RBD’s hinge with S1 (516-533) was 20% deuterated. A central helix (784-795), a turn-helix-turn (939-977) and a helix U-turn (753-759) were also moderately deuterated (28-30%). The N-terminal domain (NTD) showed an outer segment (171-176) with moderate labeling (27%).

### State difference (heat-treatment minus control)

Visualization of (heat – control) deuteration (Figure 5B), showed both increased and decreased labeling in different regions. Regions with no significant deuteration difference (within the ± 0.9 Da error of labeling estimation based upon average peptide length and repeatability, Table S2) were not interpreted to have “zero dynamic change” (25). Deuterated glycopeptide data provided coverage essential for revealing increased labeling with longer D_2_O exposure times around sequons N61 (11-15%) and N234 (11-15%) in the NTD, N343 in the RBD (10-15%), N603 (14-19%) and N801 (9%) in the S2 domain, as well as N1074 (10-16%) and N1134 (13-6%) in the stalk. Interestingly, only N1134 glycopeptides in the stalk region showed diminishing differences from the control state with increasing D_2_O exposure.

The region with lowest deuteration in the control state (trimer core interface) showed substantial increases in deuteration (764-782 helix, 21%, 1007-1024 helix, 20%, 1050-1062 β-strand, 20%). Regions with the highest deuteration in the control state showed only small increases [the “hinge” between head and stalk (1132-1145, 6%), the 630 loop (624-635, 5%)] or even reductions [the furin site (672-703, -8%) and fusion peptide (823-851, -2%)]. Data for changes in the “hinge” region were entirely from deuterated glycopeptides.

Increases in the deuteration of the RBD indicated loosening of its sub-domain architecture, including outer loop 336-350 (10-24%) and the inner β-strand 393-399 (27%). In addition, the “hinge” of the RBD to S1 (515-533) and a loop of its RBM (442-453) had lower deuteration (-4%) after heat treatment. Furthermore, core helices that appeared to be primed for fusion based on their substantial deuteration in the control state were slightly lower (939-945 and 967-977, -5%) after heat treatment. Finally, the NTD showed spike’s highest reduction in deuteration (-10%) after heat-treatment at its outermost tip (243-262). All these results are consistent with increased accessibility to limited trypsin digestion (Figure S1), looser inter-monomer interaction, and disruption of tight packing between 3 NTDs and 3 RBDs from heat treatment.

Several recent reports describe bimodality in isotopic envelopes for certain regions of spike (13, 18, 39); the only apparently bimodal peptide 878-904 in this study was interpreted to be coelution of two precursors based on overlapping isotopic envelopes even at 0 sec (data not shown).

## CONCLUSIONS

Comparing native and heat denatured SARS-CoV-2 spike demonstrated the utility of including deuterated glycopeptides in the analysis of conformational dynamics by HDX-MS. It indicated that 1) inclusion of 9 N-glycosylation sequons increased sequence coverage from 76% to 84%, 2) glycopeptides become deuterated and provide useful regional and protein-level information, 3) observed glycan identities are consistent with publications from non-deuterating studies (11, 22, 29), and 4) labeling of the amide nitrogens on N-acetyl moieties of glycans merits inclusion in estimates of glycopeptide maximum deuteration.

Inclusion of deuterated glycopeptides in HDX-MS is a step forward in attempts to measure protein dynamics under conditions as native as possible. Direct detection of deuterated glycopeptides avoids additional procedures for analyzing glycoproteins, such as deglycosylation with PNGase before or after deuteration using isoforms Rc (40), or H (41, 42). De-glycosylation adds time and complexity to sample preparation and potentially introduces artifacts during HDX-MS analyses. In the case of SARS-CoV-2 spike, reports indicate glycans participate in modulation of the RBD moving to “up” or “down” positions (43), as well as in ACE2 interaction (44, 45), indicating the importance of measuring spike glycopeptide dynamics using HDX-MS in a native state.

Several HDX-MS publications have indicated gaps in spike sequence coverage arising from “missing” glycopeptides (13, 17, 18). It is clearly advantageous to improve glycopeptide detection in HDX-MS, but this is a multi-faceted challenge at the levels of LC-MS/MS and data acquisition as well as the level of data processing. For example, HDX-MS LC is typically conducted at relatively high flow rates (40 μL/min or higher, (46)) with large-bore chromatography columns (1.0 or 2.1 mm) and steep gradients (10 to 20 minutes) to minimize deuterium back-exchange (47). All these LC conditions are non-ideal for sensitive glycopeptide detection, which performs best at nano-flow rates (< 1 μL/min) with nanoelec-trospray ion sources and columns, and long gradients (1 to 2 hours). Given these limitations of HDX-MS LC, there is likely to be considerable room for improvement in the ionization, detection, and analysis of deuterated glycopeptides.

## ASSOCIATED CONTENT

### Supporting Information

FASTA sequences of spike variants, heat denatured spike analyzed by SDS-PAGE, table of results by N-glycosylation sequon, (glyco)peptide coverage map of 614G spike, two patterns of D_2_O uptake at N61, fragment ion isotopologue analysis of glycopeptide 1132-1145 HexNAc(4)Hex(3)Fuc(1) (PDF)

The Supporting Information is available free of charge on the ACS Publications website.

## Author Contributions

D.W., T.K. and C.H. designed the study. S.O. prepared spike protein and B.M. performed heat-treatment. C.H. and T.K. performed HDX-MS. All authors contributed to designing the manuscript’s structure and C.H. wrote the manuscript. ‡These authors contributed equally.

## Notes

The authors declare no competing financial interest. The findings and conclusions in this report are those of the authors and do not necessarily represent the official position of the Centers for Disease Control and Prevention. Use of trade names and commercial sources in this presentation is for identification only and does not imply endorsement by the Division of Laboratory Sciences, National Center for Environmental Health, Centers for Disease Control and Prevention, the Public Health Service, or the U.S. Department of Health and Human Services.

## ACKNOWLEDGMENT

We thank Dr. Bin Zhou for the plasmids expressing the spike constructs used in this work.

---

“614G S6P Spike” used in HDX-MS analysis

MFVFLVLLPLVSSQCVNLTTRTQLPPAYTNSFTRGVYYPDKVFRS SVLHSTQDLFLPFFSNVTWFHAIHVSGTNGTKRFDNPVLPFND GVYFASTEKSNIIRGWIFGTTLDSKTQSLLIVNNATNVVIKVCEFQ FCNDPFLGVYYHKNNKSWMESEFRVYSSANNCTFEYVSQPFLM DLEGKQGNFKNLREFVFKNIDGYFKIYSKHTPINLVRDLPQGFSA LEPLVDLPIGINITRFQTLLALHRSYLTPGDSSSGWTAGAAAYYVG YLQPRTFLLKYNENGTITDAVDCALDPLSETKCTLKSFTVEKGIYQ TSNFRVQPTESIVRFPNITNLCPFGEVFNATRFASVYAWNRKRIS NCVADYSVLYNSASFSTFKCYGVSPTKLNDLCFTNVYADSFVIRG DEVRQIAPGQTGKIADYNYKLPDDFTGCVIAWNSNNLDSKVGG NYNYLYRLFRKSNLKPFERDISTEIYQAGSTPCNGVEGFNCYFPL QSYGFQPTNGVGYQPYRVVVLSFELLHAPATVCGPKKSTNLVKN KCVNFNFNGLTGTGVLTESNKKFLPFQQFGRDIADTTDAVRDPQ TLEILDITPCSFGGVSVITPGTNTSNQVAVLYQGVNCTEVPVAIHA DQLTPTWRVYSTGSNVFQTRAGCLIGAEHVNNSYECDIPIGAGI CASYQTQTNSPGSASSVASQSIIAYTMSLGAENSVAYSNNSIAIPT NFTISVTTEILPVSMTKTSVDCTMYICGDSTECSNLLLQYGSFCTQ LNRALTGIAVEQDKNTQEVFAQVKQIYKTPPIKDFGGFNFSQILP DPSKPSKRSPIEDLLFNKVTLADAGFIKQYGDCLGDIAARDLICAQ KFNGLTVLPPLLTDEMIAQYTSALLAGTITSGWTFGAGPALQIPF PMQMAYRFNGIGVTQNVLYENQKLIANQFNSAIGKIQDSLSSTP SALGKLQDVVNQNAQALNTLVKQLSSNFGAISSVLNDILSRLDPP EAEVQIDRLITGRLQSLQTYVTQQLIRAAEIRASANLAATKMSEC VLGQSKRVDFCGKGYHLMSFPQSAPHGVVFLHVTYVPAQEKNF TTAPAICHDGKAHFPREGVFVSNGTHWFVTQRNFYEPQIITTDN TFVSGNCDVVIGIVNNTVYDPLQPELDSFKEELDKYFKNHTSPD VDLGDISGINASVVNIQKEIDRLNEVAKNLNESLIDLQELGKYEQ GSGSGSGSGYIPEAPRDGQAYVRKDGEWVLLSTFLGSGSGSGH HHHHHGLNDIFEAQKIEWHE

“Omicron S6P Spike” used in heat-treatment and 1D gel analysis

MFVFLVLLPL VSSQCVNLTT RTQLPPAYTN SFTRGVYYPD KVFRSSVLHS TQDLFLPFFS NVTWFHVISG TNGTKRFDNP VLPFNDGVYF ASIEKSNIIR GWIFGTTLDS KTQSLLIVNN ATNVVIKVCE FQFCNDPFLD HKNNKSWMES EFRVYSSANN CTFEYVSQPF LMDLEGKQGN FKNLREFVFK NIDGYFKIYS KHTPIIVREP EDLPQGFSAL EPLVDLPIGI NITRFQTLLA LHRSYLTPGD SSSGWTAGAA AYYVGYLQPR TFLLKYNENG TITDAVDCAL DPLSETKCTL KSFTVEKGIY QTSNFRVQPT ESIVRFPNIT NLCPFDEVFN ATRFASVYAW NRKRISNCVA DYSVLYNLAP FFTFKCYGVS PTKLNDLCFT NVYADSFVIR GDEVRQIAPG QTGNIADYNY KLPDDFTGCV IAWNSNKLDS KVSGNYNYLY RLFRKSNLKP FERDISTEIY QAGNKPCNGV AGFNCYFPLR SYSFRPTYGV GHQPYRVVVL SFELLHAPAT VCGPKKSTNL VKNKCVNFNF NGLKGTGVLT ESNKKFLPFQ QFGRDIADTT DAVRDPQTLE ILDITPCSFG GVSVITPGTN TSNQVAVLYQ GVNCTEVPVA IHADQLTPTW RVYSTGSNVF QTRAGCLIGA EYVNNSYECD IPIGAGICAS YQTQTKSHGS ASSVASQSII AYTMSLGAEN SVAYSNNSIA IPTNFTISVT TEILPVSMTK TSVDCTMYIC GDSTECSNLL LQYGSFCTQL KRALTGIAVE QDKNTQEVFA QVKQIYKTPP IKYFGGFNFS QILPDPSKPS KRSPIEDLLF NKVTLADAGF IKQYGDCLGD IAARDLICAQ KFKGLTVLPP LLTDEMIAQY TSALLAGTIT SGWTFGAGPA LQIPFPMQMA YRFNGIGVTQ NVLYENQKLI ANQFNSAIGK IQDSLSSTPS ALGKLQDVVN HNAQALNTLV KQLSSKFGAI SSVLNDIFSR LDPPEAEVQI DRLITGRLQS LQTYVTQQLI RAAEIRASAN LAATKMSECV LGQSKRVDFC GKGYHLMSFP QSAPHGVVFL HVTYVPAQEK NFTTAPAICH DGKAHFPREG VFVSNGTHWF VTQRNFYEPQ IITTDNTFVS GNCDVVIGIV NNTVYDPLQP ELDSFKEELD KYFKNHTSPD VDLGDISGIN ASVVNIQKEI DRLNEVAKNL NESLIDLQEL GKYEQGSGSG SGSGYIPEAP RDGQAYVRKD GEWVLLSTFL GSGSGSGHHH HHHGLNDIFE AQKIEWHE

**Figure S1.**
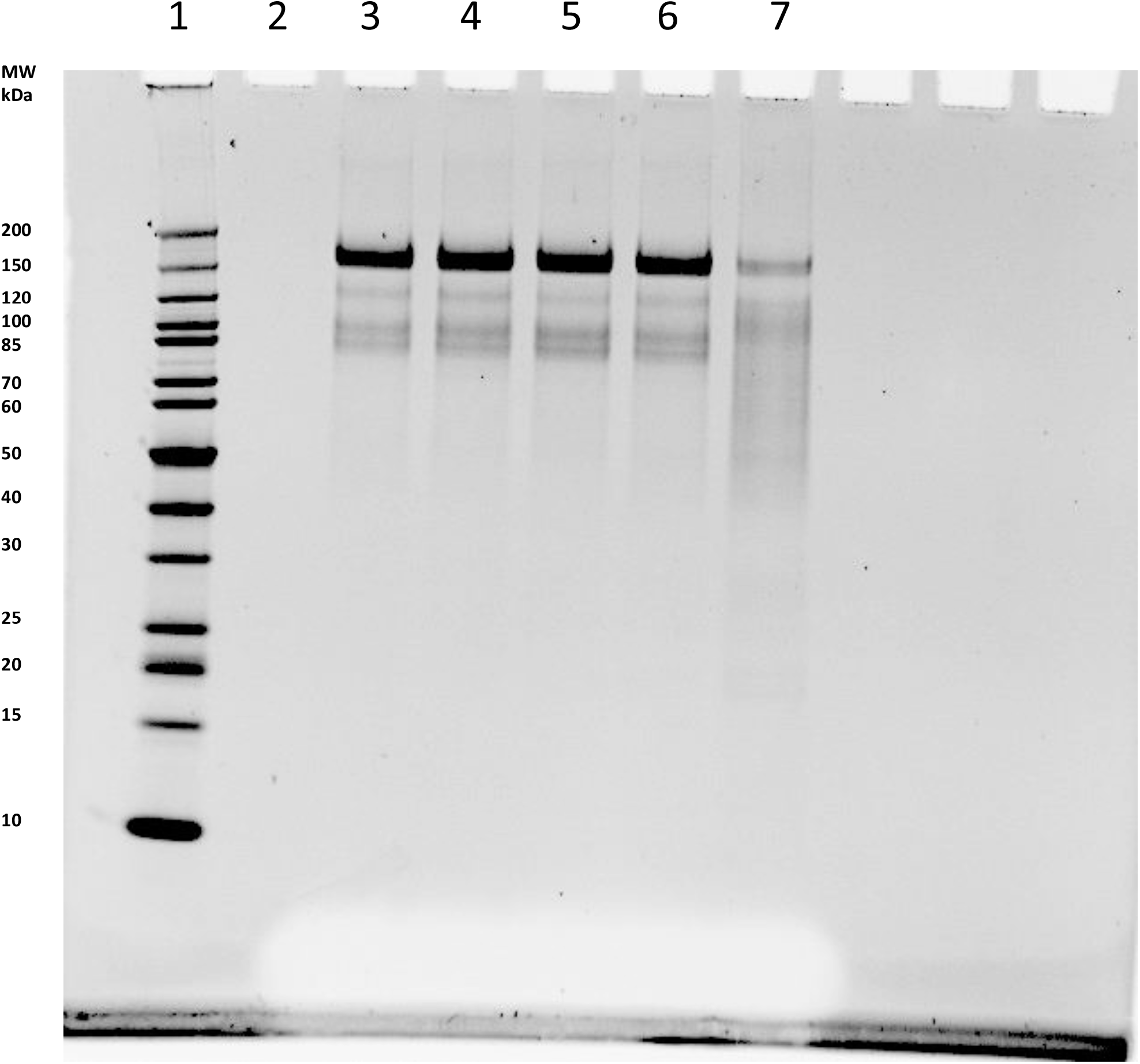
The effect of temperature and limited trypsin digestion on the conformation of SARS-CoV-2 spike protein (Omicron variant, Hexapro, His tag). Lane 1: PageRuler Unstained Protein Ladder, component MWs as indicated; Lane 2: empty; Lane 3: 0.5 μg spike, 25°C I hr, trypsin digestion as described in Methods; Lane 4: 0.5 μg spike, 35°C I hr, trypsin digestion as described in Methods; Lane 5: 0.5 μg spike, 45°C I hr, trypsin digestion as described in Methods; Lane 6: 0.5 μg spike, 55°C I hr, trypsin digestion as described in Methods; Lane 7: 0.5 μg spike, 65°C I hr, trypsin digestion as described in Methods. Gel is a non-reducing 4-12% Bis-Tris run in MES buffer at 100 V for 36 min, bands were visualized with SYPRO Ruby Protein Gel Stain (Life Technologies #S12000).

**Figure S2.**
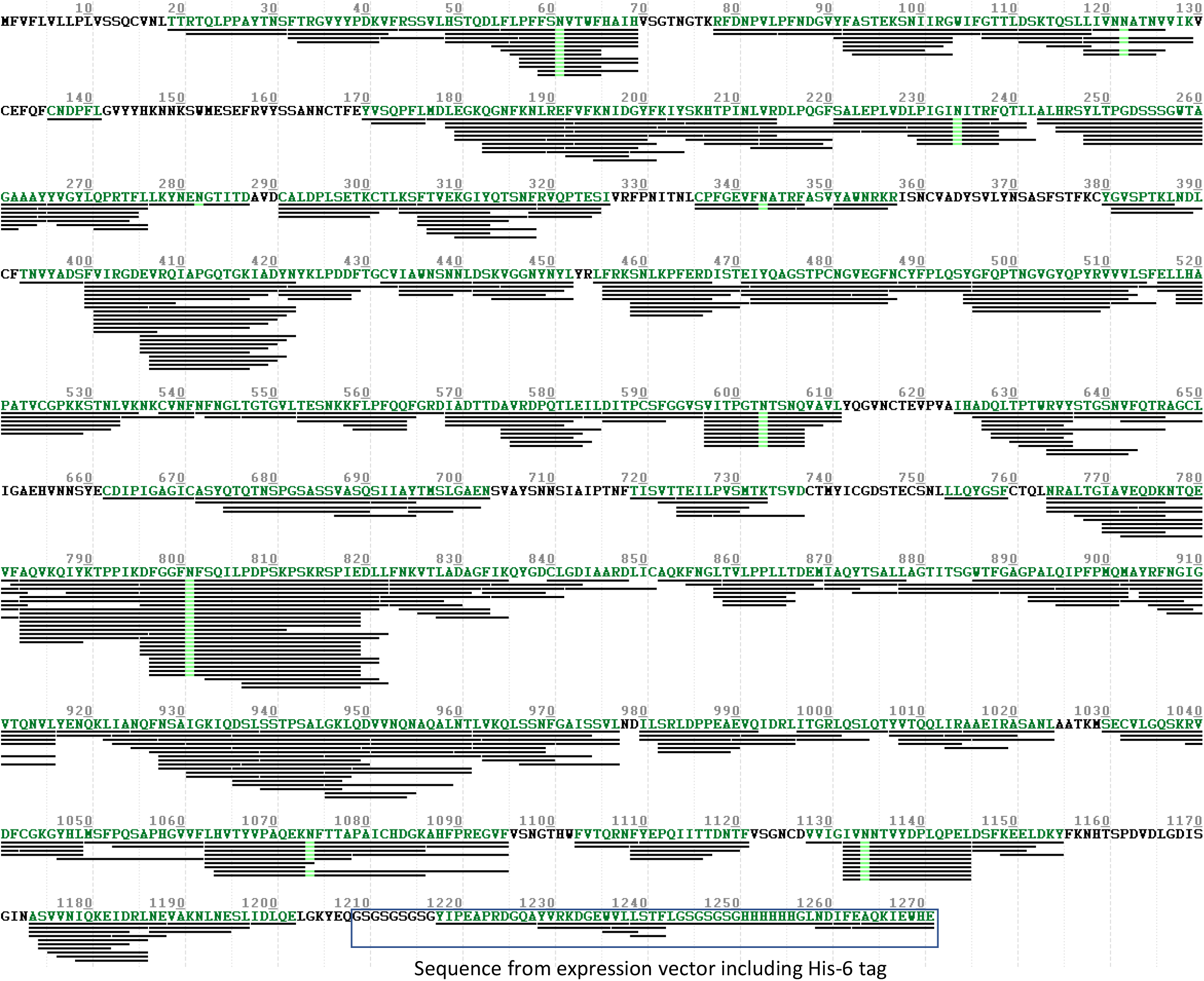
Peptide coverage map of 614G HexaPro Spike protein. Green font is protein coverage (84%), green lines under N residues indicate glycosylation detected at that N-glycosylation sequon..

**Figure S3.**
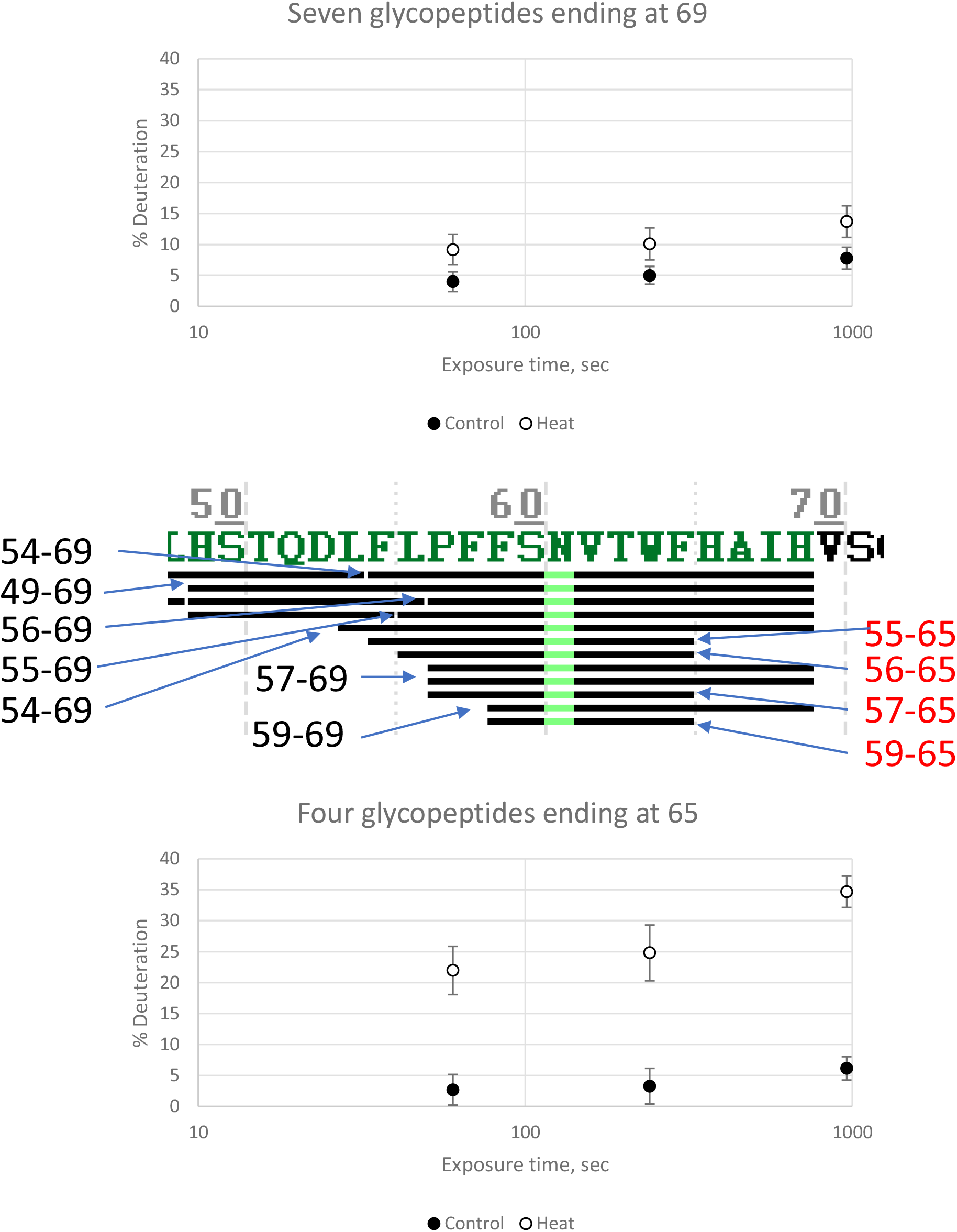
Distinct glycopeptide deuterium uptake patterns at sequon N61 of SARS-CoV-2 spike protein. Data for all glycopeptides covering N61 (middle panel) shows higher uptake for the seven glycopeptides not including sequence HAIH (lower panel) compared to the four glycopeptides including this sequence (upper panel).

**Figure S4.**
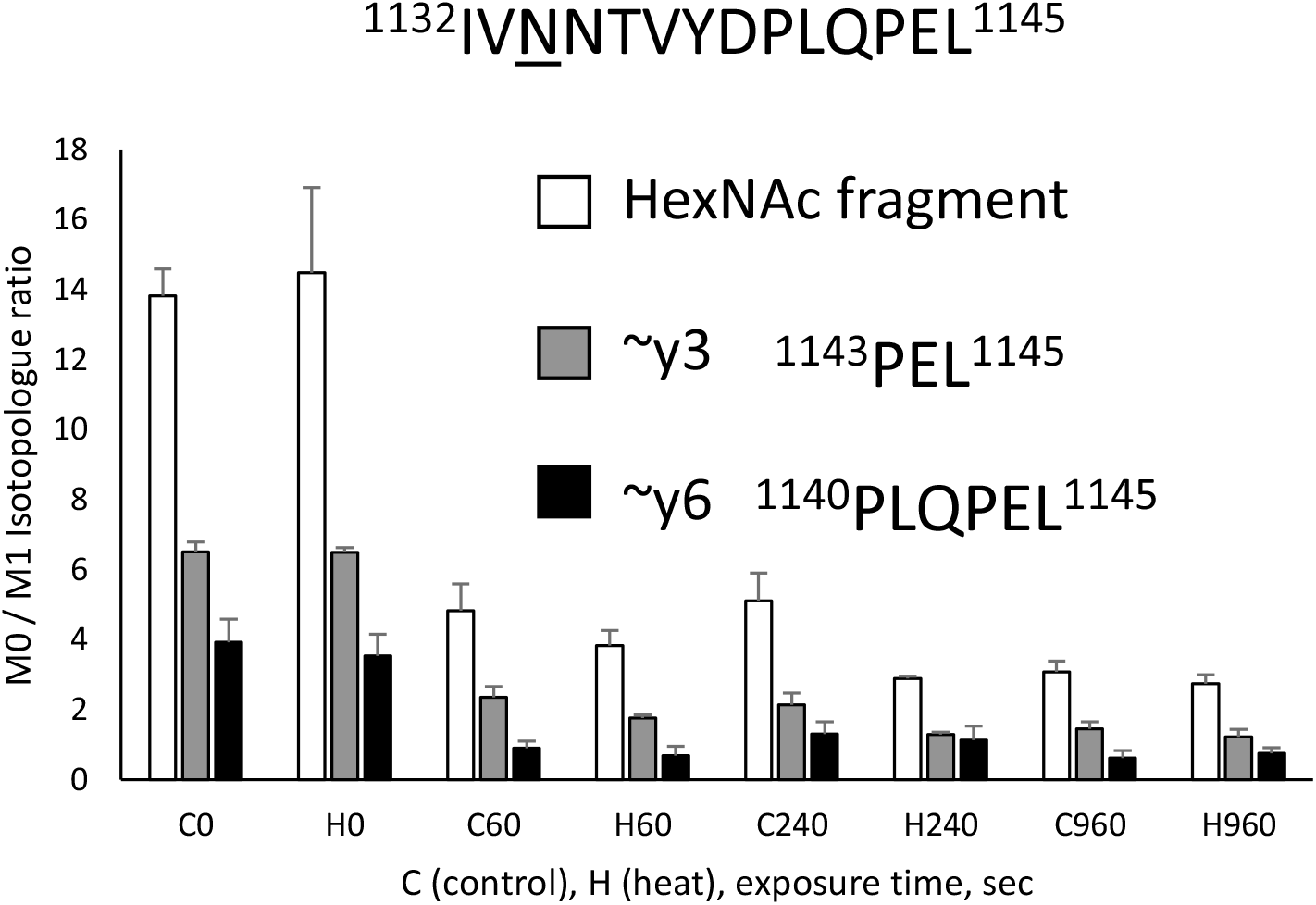
Isotopologue ratios of glycopeptide ^1132^IV<u>N</u>NTVYDPLQPEL^1145^ HexNAc(4)Hex(3)Fuc(1) fragment ions. The ratio of M0 to M1 was measured for HexNAc (204/205), ∼y3 (358/359) and ∼y6 (696/697) in 2 or 3 technical replicates at each state and exposure time. As fragment ions shift to a deuterated distribution the ratio decreases. N = 3 to 6 averaged MS2 scans for each bar.

## REFERENCES

1. Vinciauskaite V, Masson GR. Fundamentals of HDX-MS. Essays Biochem. 2023;67(2):301–14.

2. Benhaim M, Lee KK, Guttman M. Tracking Higher Order Protein Structure by Hydrogen-Deuterium Exchange Mass Spectrometry. Protein Pept Lett. 2019;26(1):16–26.

3. Oganesyan I, Lento C, Wilson DJ. Contemporary hydrogen deuterium exchange mass spectrometry. Methods. 2018;144:27–42.

4. Liu XR, Zhang MM, Gross ML. Mass Spectrometry-Based Protein Footprinting for Higher-Order Structure Analysis: Fundamentals and Applications. Chem Rev. 2020;120(10):4355–454.

5. Haque HME, Mantis NJ, Weis DD. High-Throughput Epitope Mapping by Hydrogen Exchange-Mass Spectrometry. J Am Soc Mass Spectrom. 2023;34(1):123–7.

6. Zhu S, Liuni P, Chen T, Houy C, Wilson DJ, James DA. Epitope screening using Hydrogen/Deuterium Exchange Mass Spectrometry (HDX-MS): An accelerated workflow for evaluation of lead monoclonal antibodies. Biotechnol J. 2022;17(2):e2100358.

7. Hanke L, Sheward DJ, Pankow A, Vidakovics LP, Karl V, Kim C, et al. Multivariate mining of an alpaca immune repertoire identifies potent cross-neutralizing SARS-CoV-2 nanobodies. Sci Adv. 2022;8(12):eabm0220.

8. Seow J, Khan H, Rosa A, Calvaresi V, Graham C, Pickering S, et al. A neutralizing epitope on the SD1 domain of SARS-CoV-2 spike targeted following infection and vaccination. Cell Rep. 2022;40(8):111276.

9. Zhao P, Praissman JL, Grant OC, Cai Y, Xiao T, Rosenbalm KE, et al. Virus-Receptor Interactions of Glycosylated SARS-CoV-2 Spike and Human ACE2 Receptor. Cell Host Microbe. 2020;28(4):586–601 e6.

10. Wang D, Baudys J, Bundy JL, Solano M, Keppel T, Barr JR. Comprehensive Analysis of the Glycan Complement of SARS-CoV-2 Spike Proteins Using Signature Ions-Triggered Electron-Transfer/Higher-Energy Collisional Dissociation (EThcD) Mass Spectrometry. Anal Chem. 2020;92(21):14730–9.

11. Shajahan A, Pepi L, Kumar B, Murray N, Azadi P. Site Specific N-and O-glycosylation mapping of the Spike Proteins of SARS-CoV-2 Variants of Concern. Res Sq. 2022.

12. Zhu B, Chen Z, Shen J, Xu Y, Lan R, Sun S. Structural-and Site-Specific N-Glycosylation Characterization of COVID-19 Virus Spike with StrucGP. Anal Chem. 2022;94(36):12274–9.

13. Braet SM, Buckley TSC, Venkatakrishnan V, Dam KA, Bjorkman PJ, Anand GS. Timeline of changes in spike conformational dynamics in emergent SARS-CoV-2 variants reveal progressive stabilization of trimer stalk with altered NTD dynamics. Elife. 2023;12.

14. Mehra R, Kepp KP. Structure and Mutations of SARS-CoV-2 Spike Protein: A Focused Overview. ACS Infect Dis. 2022;8(1):29–58.

15. Campos D, Girgis M, Sanda M. Site-specific glycosylation of SARS-CoV-2: Big challenges in mass spectrometry analysis. Proteomics. 2022;22(15-16):e2100322.

16. Newby ML, Fogarty CA, Allen JD, Butler J, Fadda E, Crispin M. Variations within the Glycan Shield of SARS-CoV-2 Impact Viral Spike Dynamics. J Mol Biol. 2023;435(4):167928.

17. Raghuvamsi PV, Tulsian NK, Samsudin F, Qian X, Purushotorman K, Yue G, et al. SARS-CoV-2 S protein:ACE2 interaction reveals novel allosteric targets. Elife. 2021;10.

18. Calvaresi V, Wrobel AG, Toporowska J, Hammerschmid D, Doores KJ, Bradshaw RT, et al. Structural dynamics in the evolution of SARS-CoV-2 spike glycoprotein. Nat Commun. 2023;14(1):1421.

19. Federico M. How Do Anti-SARS-CoV-2 mRNA Vaccines Protect from Severe Disease? Int J Mol Sci. 2022;23(18).

20. Alabsi S, Dhole A, Hozayen S, Chapman SA. Angiotensin-Converting Enzyme 2 Expression and Severity of SARS-CoV-2 Infection. Microorganisms. 2023;11(3).

21. Narang D, James DA, Balmer MT, Wilson DJ. Protein Footprinting, Conformational Dynamics, and Core Interface-Adjacent Neutralization “Hotspots” in the SARS-CoV-2 Spike Protein Receptor Binding Domain/Human ACE2 Interaction. J Am Soc Mass Spectrom. 2021;32(7):1593–600.

22. Wang D, Zhou B, Keppel TR, Solano M, Baudys J, Goldstein J, et al. N-glycosylation profiles of the SARS-CoV-2 spike D614G mutant and its ancestral protein characterized by advanced mass spectrometry. Sci Rep. 2021;11(1):23561.

23. Saba J, Dutta S, Hemenway E, Viner R. Increasing the productivity of glycopeptides analysis by using higher-energy collision dissociation-accurate mass-product-dependent electron transfer dissociation. Int J Proteomics. 2012;2012:560391.

24. Zhang J, Cai Y, Xiao T, Lu J, Peng H, Sterling SM, et al. Structural impact on SARS-CoV-2 spike protein by D614G substitution. Science. 2021;372(6541):525–30.

25. Masson GR, Burke JE, Ahn NG, Anand GS, Borchers C, Brier S, et al. Recommendations for performing, interpreting and reporting hydrogen deuterium exchange mass spectrometry (HDX-MS) experiments. Nat Methods. 2019;16(7):595–602.

26. Li M, Zhu W, Zheng H, Zhang J. Efficient HCD-pd-EThcD approach for N-glycan mapping of therapeutic antibodies at intact glycopeptide level. Anal Chim Acta. 2022;1189:339232.

27. Shen Y, Xiao K, Tian Z. Site-and structure-specific characterization of the human urinary N-glycoproteome with site-determining and structure-diagnostic product ions. Rapid Commun Mass Spectrom. 2021;35(1):e8952.

28. Sanda M, Benicky J, Goldman R. Low Collision Energy Fragmentation in Structure-Specific Glycoproteomics Analysis. Anal Chem. 2020;92(12):8262–7.

29. Gong Y, Qin S, Dai L, Tian Z. The glycosylation in SARS-CoV-2 and its receptor ACE2. Signal Transduct Target Ther. 2021;6(1):396.

30. Miyagi M, Nakazawa T. Determination of pKa values of individual histidine residues in proteins using mass spectrometry. Anal Chem. 2008;80(17):6481–7.

31. Tran DT, Banerjee S, Alayash AI, Crumbliss AL, Fitzgerald MC. Slow histidine H/D exchange protocol for thermodynamic analysis of protein folding and stability using mass spectrometry. Anal Chem. 2012;84(3):1653–60.

32. Guttman M, Scian M, Lee KK. Tracking hydrogen/deuterium exchange at glycan sites in glycoproteins by mass spectrometry. Anal Chem. 2011;83(19):7492–9.

33. Hatvany JB, Gallagher ES. Hydrogen/deuterium exchange for the analysis of carbohydrates. Carbohydrate research. 2023;530:108859.

34. Bai Y, Milne JS, Mayne L, Englander SW. Primary structure effects on peptide group hydrogen exchange. Proteins. 1993;17(1):75–86.

35. Chetty PS, Mayne L, Lund-Katz S, Stranz D, Englander SW, Phillips MC. Helical structure and stability in human apolipoprotein A-I by hydrogen exchange and mass spectrometry. Proc Natl Acad Sci U S A. 2009;106(45):19005–10.

36. Woo H, Park SJ, Choi YK, Park T, Tanveer M, Cao Y, et al. Developing a Fully Glycosylated Full-Length SARS-CoV-2 Spike Protein Model in a Viral Membrane. J Phys Chem B. 2020;124(33):7128–37.

37. Kumar P, Bhardwaj T, Garg N, Giri R. Microsecond simulations and CD spectroscopy reveals the intrinsically disordered nature of SARS-CoV-2 spike-C-terminal cytoplasmic tail (residues 1242-1273) in isolation. Virology. 2022;566:42–55.

38. Li F, Li W, Farzan M, Harrison SC. Structure of SARS coronavirus spike receptor-binding domain complexed with receptor. Science. 2005;309(5742):1864–8.

39. Costello SM, Shoemaker SR, Hobbs HT, Nguyen AW, Hsieh CL, Maynard JA, et al. The SARS-CoV-2 spike reversibly samples an open-trimer conformation exposing novel epitopes. Nat Struct Mol Biol. 2022;29(3):229–38.

40. Gramlich M, Maier S, Kaiser PD, Traenkle B, Wagner TR, Voglmeir J, et al. A Novel PNGase Rc for Improved Protein N-Deglycosylation in Bioanalytics and Hydrogen-Deuterium Exchange Coupled With Mass Spectrometry Epitope Mapping under Challenging Conditions. Anal Chem. 2022;94(27):9863–71.

41. Comamala G, Madsen JB, Voglmeir J, Du YM, Jensen PF, Osterlund EC, et al. Deglycosylation by the Acidic Glycosidase PNGase H(+) Enables Analysis of N-Linked Glycoproteins by Hydrogen/Deuterium Exchange Mass Spectrometry. J Am Soc Mass Spectrom. 2020;31(11):2305–12.

42. Guo RR, Comamala G, Yang HH, Gramlich M, Du YM, Wang T, et al. Discovery of Highly Active Recombinant PNGase H(+) Variants Through the Rational Exploration of Unstudied Acidobacterial Genomes. Front Bioeng Biotechnol. 2020;8:741.

43. Sztain T, Ahn SH, Bogetti AT, Casalino L, Goldsmith JA, Seitz E, et al. A glycan gate controls opening of the SARS-CoV-2 spike protein. Nat Chem. 2021;13(10):963–8.

44. Hsu YP, Frank M, Mukherjee D, Shchurik V, Makarov A, Mann BF. Structural remodeling of SARS-CoV-2 spike protein glycans reveals the regulatory roles in receptor-binding affinity. Glycobiology. 2023;33(2):126–37.

45. Casalino L, Gaieb Z, Goldsmith JA, Hjorth CK, Dommer AC, Harbison AM, et al. Beyond Shielding: The Roles of Glycans in the SARS-CoV-2 Spike Protein. ACS Cent Sci. 2020;6(10):1722–34.

46. Peterle D, DePice D, Wales TE, Engen JR. Increase the flow rate and improve hydrogen deuterium exchange mass spectrometry. J Chromatogr A. 2023;1689:463742.

47. Peterle D, Wales TE, Engen JR. Simple and Fast Maximally Deuterated Control (maxD) Preparation for Hydrogen-Deuterium Exchange Mass Spectrometry Experiments. Anal Chem. 2022;94(28):10142–50.

