## Supplemental Table 1 for "Inclusion of glycopeptides in hydrogen/deuterium exchange mass spectrometry analysis of SARS-CoV-2 spike ectodomain provides in-creased sequence coverage"

Table S1. Overview by N-glycosylation sequon of SARS-CoV-2 spike glycopeptides.

| Sequon | Glycopeptides[a] | Glycans[b] | Uptake plots consistent | Greatest $Log_2$ Fold change[c] | Lowest p-value[c] |
|---|---|---|---|---|---|
| N17 | - | - | - | - | - |
| N61 | 12 | N2H5, N2H6, N2H7 | No | 5.31 | 0.01 |
| N74 | - | - | - | - | - |
| N122 | 5 | N2H5, N2H7 | Yes | NS | NS |
| N149 | - | - | - | - | - |
| N165 | - | - | - | - | - |
| N234 | 8 | N2H9 | No | 4.40 | 0.03 |
| N282 | 1 | N2H5 | Yes | 6.61 | NS (0.07) |
| N331 | | | | | |
| N343 | 2 | N2H5 | Yes | NS | NS |
| N603 | 13 | N2H4, N2H5, N2H6, N2H7, N2H8 | Yes | 3.44 | 0.0001 |
| N616 | - | - | - | - | - |
| N657 | - | - | - | - | - |
| N709 | - | - | - | - | - |
| N717 | - | - | - | - | - |
| N801 | 27 | N2H5, N2H6, N2H7, N2H8 | No | 2.83 | 0.0004 |
| N1074 | 9 | N2H5, N2H6 | Yes | 5.04 | 0.01 |
| N1098 | - | - | - | - | - |
| N1134 | 12 | N2H5, N3H4, N3H5, N3H3F1, N3H4F1, N3H5F1, N4H3F1, N4H4F1, N3H4F1U1, N4H4F1U1 | Yes | NS | NS |
| N1158 | - | - | - | - | - |
| N1173 | - | - | - | - | - |
| N1194 | - | - | - | - | - |

a. Unique amino acid sequences, charge states and glycans are counted as different glycopeptides
b. N: HexNAc, H: Hex, F: Fuc, U: NeuAc, number indicates stoichiometry of that hexose subunit within branched glycan structure
c. Derived from volcano plot visualization of data, NS indicates not significantly different ($p > 0.05$)
